# Early mitochondrial dysfunction proceeds neuroinflammation, synaptic alteration, and autophagy impairment in hippocampus of *App* knock-in Alzheimer mouse models

**DOI:** 10.1101/2023.03.07.531542

**Authors:** Luana Naia, Makoto Shimozawa, Erika Bereczki, Xidan Li, Jianping Liu, Richeng Jiang, Nuno Santos Leal, Catarina Moreira Pinho, Erik Berger, Victoria Lim Falk, Giacomo Dentoni, Maria Ankarcrona, Per Nilsson

**Affiliations:** Department of Neurobiology, Care Sciences and Society, Division of Neurogeriatrics, Center for Alzheimer Research, Karolinska Institutet, Solna, Sweden; Department of Laboratory Medicine, Karolinska Institutet, Huddinge, Sweden; Department of Medicine, Karolinska Institutet, Huddinge, Sweden; Department of Otolaryngology Head and Neck Surgery, The First Hospital of Jilin University, Changchun, China

**Keywords:** Alzheimer’s disease, *App* knock-in mice, transcriptome analysis, mitochondria, autophagy, neuroinflammation, synaptic impairment

## Abstract

Increased amyloid β-peptide (Aβ) level is one of the drivers of Alzheimer’s disease (AD). Amyloid precursor protein (*App*) knock-in mice recapitulate the human Aβ pathology, allowing the elucidation of the downstream effects of Aβ and their temporal appearance upon disease progression. Here we have investigated the sequential onset of AD-like pathologies in the *App^NL-F^* and *App^NL-G-F^* knock-in mouse models by time-course transcriptome analysis of the hippocampus, a region severely affected in AD. Energy metabolism emerged as one of the most significantly altered pathways at an early stage of the development of the pathologies. Functional experiments in mitochondria isolated from *App^NL-G-F^* brain subsequently identified upregulation of oxidative phosphorylation driven by the activity of mitochondrial complexes I, IV and V, combined with higher susceptibility to Ca^2+^-overload. This was followed by a strong neuroinflammatory response and impaired autophagy. Accumulation of autophagosomes and reduced number of mitochondria content in presynaptic terminals could account for the altered synapse morphology including increased number of synaptic vesicles and lowered thickness of post synaptic density in *App^NL-G-F^* mice. This shows that Aβ-induced pathways in the *App* knock-in mice recapitulate some key pathologies observed in AD brain, and our data herein contributes to the understanding of their timewise appearance and potential role in new therapeutic approaches.

## INTRODUCTION

Alzheimer’s disease (AD) is the major form of dementia and leads to memory and cognitive deterioration. Brains from AD patients are characterized by disturbed protein homeostasis including extracellular deposits of amyloid β peptide (Aβ) forming plaques and intracellular accumulation of protein tau into neurofibrillary tangles ^1, 2^. The AD brain also exhibits extensive neuroinflammation and metabolic disturbances including glucose metabolism ^3^. These impairments culminate in synaptic loss and neurodegeneration which directly correlates with onset of the clinical symptoms. Thus far, only symptom-relieving medications are available. More than 400 failed clinical trials clearly showed that the underlying disease mechanisms are yet to be fully understood ^4^. The genetic association of mutations in amyloid precursor protein (APP) and presenilin 1/2, the enzymatic entities generating Aβ from APP, which induces increased production of Aβ and lead to familial AD (FAD), strongly implies Aβ as a driver of the disease ^5^. This is further supported by a protective mutation in APP which reduced Aβ levels ^6^. Aβ metabolism is also altered in the sporadic forms of AD (SAD), which comprises 99% of all AD cases, and the SAD patients display Aβ depositions in a similar spatial and temporal manner as FAD though additional genetic and environmental factors may contribute to onset and disease progression ^7, 8^. To unravel Aβ-induced pathophysiological pathways, animal models that recapitulate the Aβ pathology are powerful tools. *App* knock-in mice exhibit high levels of Aβ42 due to the Swedish (KM670/671NL) ^9^ and Beyreuther (I716F) ^10, 11^ mutations inserted in the mouse *App* gene which specifically lead to the generation of AD-causing Aβ42 species, whereas the App levels are physiological thereby circumventing potential artefacts caused by APP overexpression paradigms applied in APP transgenic mice ^12, 13^. These mutations induce Aβ plaque pathology, which starts from nine month-of-age in the *App^NL-F^* mice, leading to neuroinflammation and synaptic loss around the Aβ plaques, subsequently inducing memory impairment at 18 months-of-age. To accelerate the Aβ pathology and its downstream effects, the Arctic (E693G) mutation was additionally inserted in the mouse *App* gene in the *App^NL-G-F^* mice to induce Aβ oligomerization and protofibril formation. *App^NL-G-F^* mice show earlier onset of Aβ plaque deposition starting from two months-of-age, memory impairment from six months-of-age, and a more pronounced neuroinflammation. To understand the temporal-dependent development of pathologies associated with disease progression in hippocampus, a major area affected in AD, we performed RNA sequencing of hippocampi from both *App^NL-F^* and *App^NL-G-F^* mice at three different ages before onset, after onset and at late stage of the development of AD-like symptoms. Based on the transcriptome data we designed functional assays and histopathological examinations to assess the affected pathways. We identified early and substantial changes in mitochondrial function, followed by a strong neuroinflammatory response, declining mitochondrial function, inhibited autophagy and ultimately synaptic impairment.

## MATERIALS AND METHODS

### Animals and ethical permits

Homozygous *App^NL-F/NL-F^* (denoted herein *App^NL-F^*) and *App^NL-G-F/NL-G-F^* (denoted herein *App^NL-G-F^*) and *App^wt/wt^* (WT) (all on C57BL/6J background) were housed at Karolinska Institutet animal facility under conditions of controlled temperature (22–23 °C) and under a 12-h light/12-h dark cycle. Food and water were available *ad libitum*. All experimental procedures were carried out in accordance with the guidelines of the Institutional Animal Care and Use of Committee and the European Community directive (2010/63/EU) and approved by Linköping Animal Ethical Committee (ID 407) and Stockholm Animal Ethical Committee (15758-2019). For this study two-month-old, six-month-old, 10-12-month-old, 18-month-old and 22-to 24-month-old females were used.

### RNA isolation and sequencing

RNA was extracted from dissected hippocampal brain tissue (n = 3 females/genotype and age) preserved in RNAlater (AM7020, Thermo Scientific) using RNeasy Lipid Tissue Mini Kit (74804, Qiagen) according to manufacturer’s instructions. RNA quality (RNA integrity number, RIN) and quantity was measured in a Bioanalyzer 2100 (Agilent) using the Agilent RNA 6000 Nano Kit (part number 5067-1511). NEBNext Ultra II Directional RNA Library Prep Kit for Illumina (E7760S, New England Biolabs) was used to prepare the sequencing libraries, starting with 200 ng of total RNA. In short, mRNA was isolated and fragmented using the NEBNext poly(A) mRNA magnetic isolation module (E7490S, New England Biolabs). First and second strands of cDNA were synthesized and purified using AmPure XP beads (A63880, Beckman Coulter). Adaptor ligation and size selection were performed according to the manufacturer’s protocol. Adaptor ligated cDNA was PCR enriched to incorporate an Illumina compatible index sequence (NEBNext Multiplex Oligos for Illumina, Dual Index Primers Set1, E7600S, New England Biolabs). The libraries were purified using AmPure XP beads, the size distribution of the libraries was analyzed with the Bioanalyzer 2100 using the Agilent High Sensitivity DNA Kit (part number 5067-4626). Quantification of the libraries was performed with the Qubit® 2.0 Fluorometer (ThermoFisher Scientific) and Qubit^TM^ dsDNA HS Assay Kits (Q32851, Invitrogen). Finally, all 30 libraries were pooled and diluted to 3.5 nM for sequencing on one lane of a Hiseq 3000 sequencer (Illumina), using a single read 50 bp and dual indexed sequencing strategy.

### Sequence processing and differential gene expression analysis

All raw sequence reads available in FastQ format was mapped to the mouse genome (mm10) using Tophat2 with Bowtie2 option ^14, 15^, where adaptor sequences were removed using trim galore before read mapping. BAM files containing the alignment results were sorted according to the mapping positions. Raw read counts for each gene were calculated using featureCounts from Subread package ^16^. DEseq2 was used to perform the analysis of differential gene expression, where genes with raw counts was used as input ^17^. The differentially expressed genes (DEGs) were identified by adjusted *p* value for multiple testing using Benjamini-Hochberg correction with False Discovery Rate (FDR) values less than 0.1.

### Gene ontology enrichment analysis

The significantly DEGs were applied to Gene ontology (GO) enrichment analysis using online software AmiGO website (http://amigo.geneontology.org/amigo), and the significant enrichment GO terms was identified using Fisher’s Exact test with Bonferroni corrected *p*-values ≤ 0.05. Identification of genes involved in mitochondrial function, neuroinflammation and autophagy was performed by using free online databases and software including Uniprot (http://uniprot.org), DAVID v6.8 (https://david-d.ncifcrf.gov/) and PANTHER v13.1 (www.pantherdb.org). Genes included within the GO terms (in all ontologies: biological processes, molecular function, cellular component) containing the word “autophag” were considered genes involved in the autophagosomal-lysosomal system, those containing the word “inflamm” were considered inflammation related genes, those containing the word “mitochondri” were considered to be genes related to mitochondrial processes, and used for further enrichment analysis.

### Pathway analysis

Gene Set Enrichment Analysis (GSEA) ^18^ was applied to perform pathway analysis using the KEGG pathways dataset. First, genes were ranked decreasingly according to the Log_2_ Fold Change (Log2FC) of expression. For each query pathway, if gene *i* is a member of the pathway, it is defined as

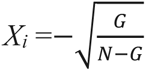

If gene *i* is not a member of the pathway, it is defined as

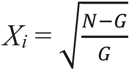

where *N* indicates the total number of genes and *G* indicates the number of genes in the query pathway. Next, a max running sum across all *N* genes Maximum Estimate Score (MES) is calculated as

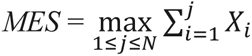

The permutation test was performed 1000 times to judge the significance of MES values, where the null hypothesis is that the pathway is not enriched in ranking. If the query pathway with a nominal *p*-value less than 0.05 and adjusted *p*-value for multiple testing using Benjamini-Hochberg correction with FDR values less than 0.1, the null hypothesis would be rejected, and the query pathway would be considered as significantly enriched. MES value represents the expression direction of a pathway, where a positive MES value indicates up-enrichment (up-regulation) whereas a negative MES value indicates down-enrichment (down-regulation) of a pathway.

### Unsupervised genome-wide clustering

The raw read matrix distributed column-wisely with samples names and row-wisely with gene names, was normalized using between-samples normalization method implemented in DESeq2 package ^17^. Unsupervised genome-wide clustering was then performed using *t*-SNE plot with the R package Rtsne ^19^. The *t-*SNE plot is based on the 50 most variant dimensions of the initial PCA plot, wherein genes with duplicate reads are filtered out. The Speed/accuracy trade-off was set as 0.0 to exact *t-*SNE distance matrix. The perplexity is adjusted accordingly for the optimal clusters shape. Plots showing all samples are based on the *t*-SNE field parameters V1 and V2.

### cDNA reverse transcription (RT) and qPCR

100 ng of RNA was reverse transcribed using the High-Capacity cDNA Reverse Transcription Kit (Thermo Fisher Sci., Cat. no: 4374966) according to manufacturer’s instructions. The TaqMan Fast Advanced Master Mix (Thermo Fisher Sci., Cat. no: 4444557) was used to perform the qPCR using the following TaqMan mouse gene expression assays (FAM) (Thermo Fisher Sci., Cat. no: 4331182): Mm04225236_g1 for *Atp8*; Mm00432648_m1 for *Cox8b*; Mm00444593_m1 for *Ndufa1*; Mm01352366_m1 for *Sdha*; Mm01615741_gH for *Uqcrb*; Mm01183349_m1 for *Clec7a*; Mm00441259_g1 for *Ccl3*; Mm04209424_g1 for *Trem2*; Mm00437893_g1 for *C4b*; Mm01312230_m1 for *Chrna4*; Mm00495267_m1 for *Lamp2*; Mm00523599_g1 for *Rab7b*; Mm01310727 for *Rubcnl*; Mm01274264_m1 for *Trim30a*; Mn00450314_ml for *Vamp8*. Experiments were performed in the 7500 Fast Real-Time PCR System (Applied Biosystems) and each sample was run in triplicates. Gene expression was normalized to *b3-tubulin* mRNA (Mm00727586_s1).

### Brain homogenates

Right hippocampi from the same individuals used for RNA-seq experiments (*n* = 3/genotype and age) were cut in pieces while kept cold on ice and homogenized with a 2 cm^3^ glass-Teflon homogenizer in RIPA buffer (150 mM NaCl, 50 mM Tris, 1% Triton X-100, 0.5% DOC, 0.1% SDS; pH=7.5) supplemented with 1:100 proteases inhibitors (G-Biosciences, Cat. no: 786-433) and 1:100 phosphatases inhibitors (Sigma-Adrich, Cat. no: P0044) in a proportion of 1:15 (mg tissue/µl buffer). The homogenates were then sonicated for 15 sec on ice and centrifuged at 20,800 *g* for 30 min, at 4°C, to remove cell debris. Supernatant was collected and protein concentration determined by Pierce™ BCA Protein Assay Kit (Thermo Fisher Sci., Cat. no: 23225).

### Crude synaptosomal fractions

Crude synaptosomal fractions were isolated from dissected left hippocampal brain tissues (*n* = 4/genotype and age) by homogenizing the tissue with a 2 cm^3^ glass-Teflon homogenizer at 800 rpm in lysis buffer (0.32 M sucrose, 5 mM Hepes, and 10 ml ddH_2_O) supplemented with 1:100 diluted proteases inhibitors (G-Biosciences, Cat. no: 786-433) and 1:100 diluted phosphatases inhibitors (Sigma-Adrich, Cat. no: P0044) in a proportion of 1:10 (mg tissue/µl buffer). The homogenates were centrifuged at 1,000 *g* at 4 °C for 10 min to remove nuclei and debris, and the supernatant was centrifuged again at 12,000 *g* at 4 °C for 20 min. Supernatant composed of the light membrane fraction and soluble enzymes (S2) was transferred to a new tube, and the pellet containing crude synaptosomes and mitochondria (P2) was resuspended in 75 μl RIPA buffer supplemented with 1:100 diluted proteases inhibitors and 1:100 diluted phosphatases inhibitors. The protein concentration was determined by Pierce™ BCA Protein Assay Kit (Thermo Fisher Sci., Cat. no: 23225).

### Western blotting

15-20 µg of protein were loaded onto 4-12% Bis-Tris gels (Novex, Cat. no: NP0336BOX) or 4-20% Tris-Glycine extended gels (BIO-RAD, Cat. no: 4561095) for separation and transferred to nitrocellulose or methanol activated PVDF membranes. Membranes were blocked in 5% BSA TBS-T or 5% skim milk TBS-T and incubated overnight at 4°C with the following primary antibodies diluted in 5 % BSA or 5% skim milk dissolved in TBS-T: pSer293-PDH (1:1,000) (Merck Millipore, Cat. no. ABS204), PDH (1:1,000) (Santa Cruz, Cat. no. sc-377092), IDE (1:900) (Abcam, Cat. no. ab32216), LC3 (1:500) (MBL, Cat. no. PM036), pSer403-p62/SQSTM1 (1:500) (Merck Millipore, Cat. no. MABC186-I), p62/SQSTM1 (1:1.000) (Cell Signaling, Cat. no. 5114), Synaptophysin (1:500) (Merck Millipore, Cat. no. MAB5258), PSD95 (1:1,000) (Cell Signaling, Cat. no. 3409), ULK1 (1:500) (Cell signaling, Cat. no. 8054), pSer555-ULK1 (1:500) (Cell Signaling, Cat. no. 5869), pSer757-ULK1 (1:500) (Cell Signaling, Cat. no. 14202), β3 tubulin (1:2,000) (Santa cruz, Cat. no. sc-80016). After washing with TBS-T for three times, membranes were incubated with fluorescent secondary antibodies (1:10,000-20,000) at room temperature for 1 h. Blots were developed using Odyssey CLx (LI-COR) and the resulting band intensities were quantified by using Image Studio Lite (LI-COR).

### Immunofluorescence staining

Brains were dissected from two- and 12-month-old WT and *App^NL-G-F^* mice (*n* = 4/genotype) after perfusion with PBS under anesthetization by isoflurane, and fixed in 10% formalin solution (Merck Millipore, Cat. no: HT501128). Brains were dehydrated and the paraffin-embedded brains were sliced into 4 μm sections by microtome (Microm HM 360). For triple staining, after deparaffinization and antigen retrieval, sections were stained using Opal 4-Color IHC kit (Akoya, NEL810001KT) according to manufacturer’s instructions with the following primary antibodies: LC3A and B (1:2,000) (Novus, Cat. no. NB100-2220), synaptophysin (1:400) (Synaptic Systems, Cat. no. 101 002). After LC3 and synaptophysin staining, sections were incubated overnight at 4°C with anti-Aβ antibody 82E1 (1:1,000) (IBL, Cat. no. 10323) diluted in PerkinElmer antibody diluent/blocking buffer, followed by incubation with anti-mouse IgG antibody conjugated with Alexa Fluor 647 (1:200) (Thermo Fisher Scientific, Cat. no. A-21235) diluted in PerkinElmer antibody diluent/blocking buffer at room temperature for 1 h. For double staining, after deparaffinization and antigen retrieval, sections were blocked with 5% normal goat serum in PBS-T. Sections were incubated overnight at 4°C with the primary antibody for LC3A and B (1:2,000) (Novus, Cat. no. NB100-2220) diluted in 5% normal goat serum in PBS-T. After washing, sections were incubated with biotinylated anti-rabbit IgG antibody (1:200) (Vector, Cat. no. BA-1000) diluted in 5% normal goat serum in PBS-T for 1 h at room temperature. The signals were amplified by TSA Fluorescein system (Akoya Biosciences, Cat. no. NEL701A001KT) according to manufacturer’s instructions. Sections were incubated overnight at 4°C with anti-Aβ antibody 82E1 (1:1,000) (IBL, Cat. no. 10323) diluted in 5% normal goat serum in PBS-T, followed by incubation with Alexa Fluor 546 conjugated anti-mouse IgG antibody (1:200) (Thermo Fisher Scientific, Cat. no. A-11030) diluted in 5% normal goat serum in PBS-T for 1 h at room temperature. Nuclei were counterstained with Hoechst 33342 (1:1,000). The images were acquired using Zeiss LSM980 and Nikon fluorescence microscope (Nikon, eclipse E800).

### Isolation of functional mitochondria

Hippocampi were dissected from mouse brains (*n* = 5 – 6/genotype and age) and washed once in ice-cold PBS. Mitochondria were isolated using discontinuous Percoll density gradient centrifugation as previously described ^20^. Briefly, hippocampi were transferred to an 8 ml Dounce tissue grinder of the Potter-Elvehjem PTFE and homogenized approximately 6 ties in ice-cold mitochondria isolation buffer (225 mM mannitol, 75 mM sucrose, 1 mM EGTA, 5 mM HEPES–KOH, pH 7.2) supplemented with 1 mg/ml fatty acid-free BSA. The final homogenate was centrifuged at 1,100 *g* at 4°C for 2 min. The supernatant was collected and mixed with fresh 80% Percoll (GE Healthcare, Cat. no. 17-5445-02) prepared in mitochondrial dilution buffer (1000 mM sucrose, 50 mM HEPES– KOH, 10 mM EGTA, pH 7.0), to obtain a 5% Percoll solution, which was further carefully layered on the top of the fresh 10% Percoll. The mitochondrial fraction was pelleted by centrifugation at 18,500 *g* at 4°C for 10 min. The pellet was then resuspended in 1 ml of mitochondria washing buffer (250 mM sucrose, 5 mM HEPES–KOH, 0.1 mM EGTA, pH 7.2) and centrifuged at 10,000 *g* at 4°C for 5 min. Mitochondrial pellet was again resuspended in a small volume of ice cold MWB, to concentrate mitochondria that was kept in ice for further analysis for a maximum of 3 hours. Protein content of isolated mitochondria was quantified using Pierce™ BCA Protein Assay Kit.

### Oxygen consumption rate (OCR) evaluation

2.5 µg of isolated mitochondria diluted in mitochondrial assay solution (70 mM sucrose, 220 mM mannitol, 10 mM K_2_HPO_4_, 5 mM MgCl_2_, 1 mM EGTA, 2 mM HEPES–KOH) supplemented with 0.2% (w/v) fatty acid-free BSA were seeded in poly(ethyleneimine)-coated (1:15,000; Sigma-Aldrich, Cat. no: 03880) XFe96 seahorse plates by centrifugation at 2,000 g for 18 min at 4°C. For respiratory coupling, MAS was further supplemented with 10 mM succinate plus 2 μM rotenone to ensure state II respiration, followed by the injection of 4 mM ADP to induce state III, which was then inhibited by addition of 3.2 μM oligomycin. The addition of 4 μM uncoupler FCCP (state IIIu) reflects the maximal respiratory chain activity as well as the maximal substrate oxidation rate. Finally, 4 μM antimycin A was added to fully block the respiratory chain and the residual OCR. To determine electron flow activity, *i.e*., sequential determination of complexes I-IV-dependent respiration, MAS was supplemented with 10 mM pyruvate, 2 mM malate and 4 µM FCCP. Sequential injection of rotenone (2 μM; complex I inhibitor), succinate (10 mM; complex II substrate), antimycin A (4 μM; complex III inhibitor) and ascorbate/TMPD (N,N,N′,N′-tetramethyl p-phenylenediamine) (10 mM/100 μM; electron donors to cytochrome C/complex IV) were performed to evaluate individual mitochondrial complexes activity ^20, 21^.

### Calcium retention capacity

Calcium (Ca^2+^) uptake by isolated mitochondria was measured using the Ca^2+^ sensitive probe Calcium Green-5N. Briefly, 10 μg of mitochondria were incubated in mitochondrial reaction buffer (100 mM sucrose, 100 mM KCl, 2 mM KH_2_PO_4_, 5 mM HEPES, 0.01 mM EGTA, 3 mM succinate, 3 mM glutamate, 0.1 mM ADP-K, pH 7.4) supplemented with 1 μM oligomycin and 150 nM Calcium Green-5N (Thermo Fisher Sci., Cat. no: C3737) ^20, 22^. Mitochondria were then dispensed in a 96-multiwell plate and fluorescence was measured in the microplate reader Fluostar Galaxy (LabVision) by excitation at 506 nm and emission at 523 nm. After a baseline of 3 min, pulses of 10 μM CaCl_2_ were added to mitochondria in 3 min intervals. Mitochondrial Ca^2+^ retention capacity was calculated by the area under the curve after CaCl_2_ pulses, which indicates the amount of extramitochondrial Ca^2+^ taken up by mitochondria.

### Transmission electron microscopy

Ten- to twelve-month-old *App^NL-G-F^* and 22- to 24-month-old *App^NL-F^* mice were anaesthetized and perfused through intracardial perfusion with 2% glutaraldehyde and 1% formaldehyde in 0.1 M phosphate buffer (n = 4/genotype). Hemispheres were cut in a brain slicer matrix and coronal slices collected for sectioning. Leica Ultracut UCT or EM UC7 was used to create ultrathin sections, with uranyl acetate and lead citrate used as contrasting agents. For mitochondria and endoplasmic reticulum (ER) analysis, sections were examined at 100 kV using a Tecnai 12 BioTWIN transmission electron microscope. Images were acquired from the hippocampus *Cornu Ammonis* area 1 (CA1) at a primary magnification of 30,000 ×. Seven different cells were snapped per brain and 30-40 synapses were analyzed per condition. A synapse was considered when the presence of both synaptic vesicles and post-synaptic density were detected. Synaptic vesicles were counted using the cell counter plugin from ImageJ. ER aspect ratio was quantified by dividing the major axis of the ER profile to the minor axis. For the observation of autophagic vesicle accumulation, images were acquired from the hippocampus CA1 at a primary magnification of 16,500 × or 26,500 ×.

### CSF sampling and proximity extension assay (PEA)

Mice were anesthetized with 1.5 % isoflurane and thereafter placed in a stereotactic instrument with 120-130° head to body angle (n = 5/genotype). Sagittal incision was made posterior to the occipital crest. The dura mater was exposed by a gentle blunt dissection of subcutaneous tissue and muscles under dissection microscope and cleaned by cotton swabs soaked in PBS to remove any blood contamination. The dura mater was punctured with a 27-gauge needle, avoiding visible blood vessels, and CSF was subsequently collected in a glass capillary tube. Collected CSF was inspected under the microscope and discarded in the case of detected blood contamination. Samples were collected in low-affinity polypropylene tubes, snap-frozen in liquid nitrogen and stored at – 80 °C until further analysis. Levels of 92 proteins in CSF were analyzed using Target 96 Mouse Exploratory Panel with PEA technology, according to manufacturer’s instructions (Olink Proteomics).

### Statistical Analysis

Statistical analysis was done by using GraphPad Prism version 9 and data were presented as mean ± SEM. n refers to the number of unique individual biological samples. Non-parametric Kruskal-Wallis tests followed by Dunn’s multiple comparison test or unpaired *t*-test (two tailed) were performed as indicated in the individual figure legends. Outliers were detected using ROUT method (Q=1%).

## RESULTS

### Altered transcriptomes in *App^NL-F^* and *App^NL-G-F^* mouse hippocampus

To gain a comprehensive understanding of how the progression of AD related pathologies driven by Aβ amyloidosis in the two *App* knock-in mouse models affect gene expression, we have performed a time course mRNA expression profiling of hippocampi of *App^NL-F^* and *App^NL-G-F^* mice and age matched wild type (WT) controls. *App^NL-F^* mice exhibit initial Aβ deposition at six-month-of-age and memory impairment at 18-months-of-age ^23, 24^ whereas *App^NL-G-F^* mice exhibit aggressive and early Aβ pathology with initial deposition at two months of age, while memory impairment is observed as early as six months of age ^23, 25^. This study was designed to enable detection of early alterations in hippocampus (at two and six month-of-age in *App^NL-G-F^* and *App^NL-F^* mice, respectively), pathophysiological changes associated with an established amyloidosis (at six and 12-month-of-age in *App^NL-G-F^* and *App^NL-F^* mice, respectively) as well as changes in the more advanced stages associated with memory impairment (at 12- and 18-month-of-age in *App^NL-G-F^* and *App^NL-F^* mice, respectively) by RNA sequencing (RNA-seq) analyses (**Figure 1A**). The RNA-seq results were subsequently confirmed by downstream biochemical, functional, and structural analyses as depicted in **Figure 1A**. In total 46,000 transcripts were obtained. The overview of the transcriptomes through clustering and *t*-SNE plots reveals a separation of transcriptomes of WT mice from the transcriptomes of *App* knock-in mice, whereas 18-month-old *App^NL-F^* and six-, 12-month-old *App^NL-G-F^* mice clustered together (**Figure 1B**). Notably, the number of differentially expressed genes (DEGs) is much higher in the *App^NL-G-F^* mice, especially at six and 12 months of age, as compared to the transcriptome alterations in the *App^NL-F^* mice (**Figure 1C**). In *App^NL-F^* mice, the highest number of DEGs were observed at six months of age, prior to onset of Aβ plaque pathology, and after onset of Aβ plaque pathology at 18 months of age (**Figure 1C**). In the *App^NL-G-F^* mice over 600 genes are altered as early as at two-month-of-age, and the number further increased in six months old mice, reaching over 2600 DEGs. Around 1500 DEGs were observed in the 12-month-old mice, though many of the altered genes at this age were already altered at six months of age. Volcano plots of the transcriptomes from the different cohorts further highlighted the drastic increase in DEGs in the *App^NL-G-F^* mice, especially at six and 12 months-of-age (**Figure 1D**). Many of the significantly altered genes in the *App* knock-in mice are involved in neuroinflammation (e.g., *Trem2, Ccl3* and *C4b*) and lipid and energy metabolism (e.g., *ApoE, mt-Atp8* and *Cox8b*). The transcriptomes were next subjected to Gene Set Enrichment Analysis (**Supplemental Figure 1**) which further confirmed significant alterations in pathways associated with energy production including TCA cycle and oxidative phosphorylation (OxPHOS) (**Figure 1E**). Other affected pathways included protein homeostasis and inflammation **(Figure 1E, Supplemental Figure 1)**. Notably, the “Alzheimer’s disease” pathway (containing genes related to energy metabolism, glutamate receptor signaling and apoptosis, see **Supplemental Table 1**) was upregulated in early stages and downregulated in the later stages. The “Alzheimer’s disease” pathway also contains processes such as Aβ metabolism including the Aβ degrading enzyme insulin-degrading enzyme (IDE). Biochemical analysis consistently revealed a significant decrease of IDE at both the RNA and protein levels in the hippocampus of *App^NL-F^* mice (**Figure 1F**).

**Figure 1.**
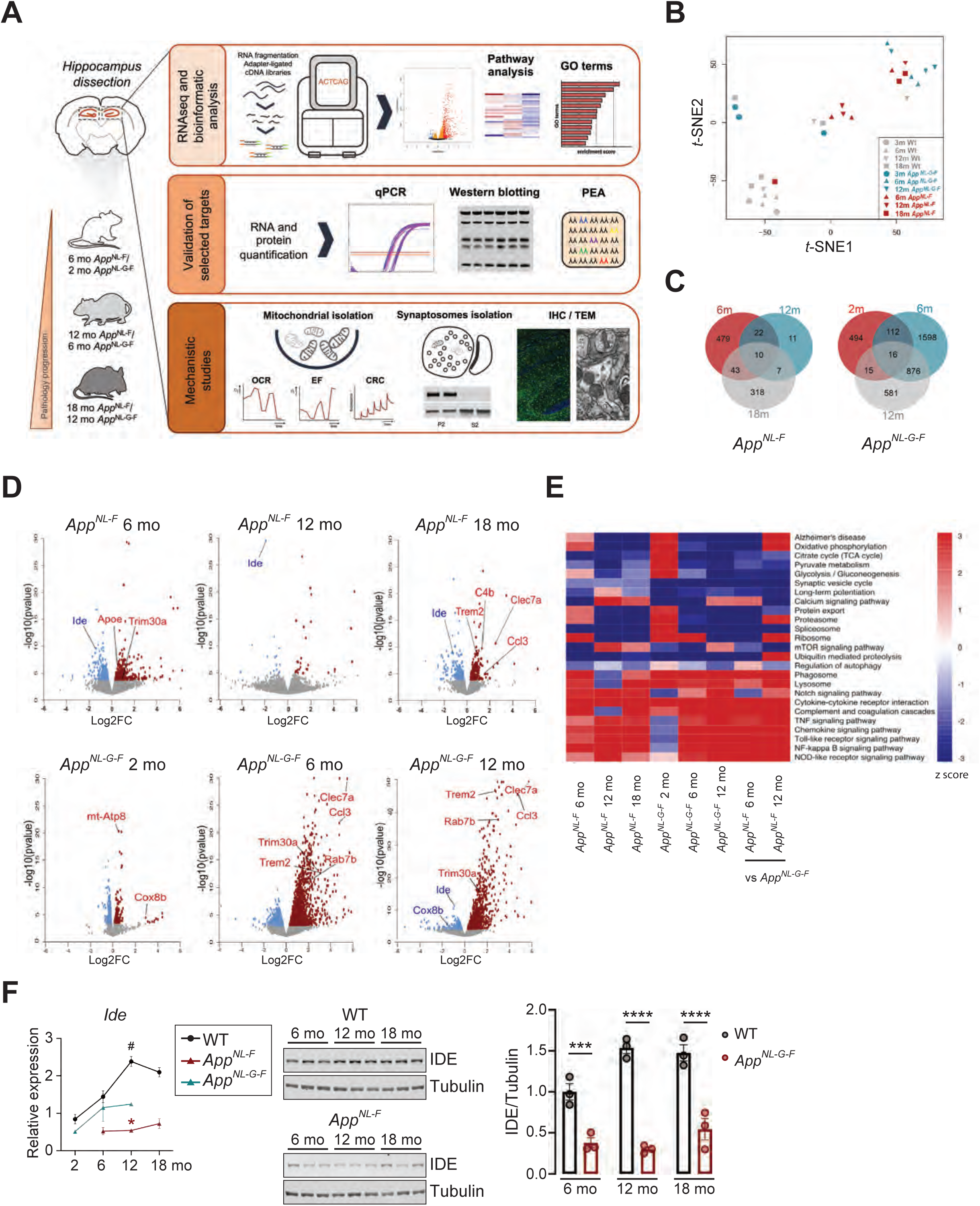
Transcriptome profiling identified Aβ-induced alterations in genes and pathways in hippocampus of *App* knock-in mice. (A) Hippocampi were dissected from two-, six-, 12-month-old *App^NL-G-F^* mice and six-, 12-, 18-month-old *App^NL-F^* mice and age matched WT controls. RNA was extracted from dissected hippocampi and cDNA libraries were synthesized. Finally, RNA sequencing was performed by a Hiseq 3000 sequencer (n = 3). Validation studies were performed by RT-qPCR using the same RNA samples for RNA sequencing. Western blotting with hippocampal brain homogenate and Olink proteomics were conducted for validations at protein level. Mitochondria and crude synaptosomal fraction were isolated for mechanistic studies of mitochondrial and autophagic functions. Mitochondrial dysfunction and autophagic alterations were revealed in *App^NL-G-F^* mice by electron microscopy. (B) *t-*SNE plot representing difference of *App^NL-F^*, *App^NL-G-F^* and WT transcriptomes. Each symbol represents one mouse individual. Each color represents one mouse genotype, red: *App^NL-F^*, blue: *App^NL-G-F^*, gray: WT, and each shape of symbol represents mouse age, circle: two-month-old, triangle: six-month-old, inverted triangle: 12-month-old, square: 18-month-old. (C) Venn diagram of significantly DEGs in *App^NL-F^* and *App^NL-G-F^* vs WT mice (FDR < 0.1). (D) Volcano plots of the gene expression profiles in different time points of *App^NL-F^* or *App^NL-G-F^* vs WT mice. Red and blue dots indicate significantly up- and downregulated genes (FDR < 0.1), respectively, in *App* knock-in mice. Grey dots indicate non-significantly altered genes. Genes validated by RT-qPCR are highlighted. (E) Heatmap of selected pathways related to AD, glucose metabolism, neuroinflammation and autophagy in *App^NL-F^* vs WT mice (columns 1 to 3), *App^NL-G-F^* vs WT mice (columns 4 to 6) and *App^NL-F^* vs *App^NL-G-F^* mice (columns 7 and 8). The *p*-value of each enriched pathway was converted to Z score. Significantly up- and downregulated pathways have the absolute value of Z scores ≥ 1.96. (F) Relative mRNA expression of *Ide* gene was normalized to *Tubb3* (n = 3, left). Statistical significance was analyzed using Kruskal-Wallis tests followed by Dunn’s multiple comparison test. \**p* < 0.05. * *App^NL-F^* mice vs age matched WT controls, ^#^ vs two-month-old WT mice. IDE protein levels in hippocampal homogenates were quantified by Western blotting and normalized to β3-tubulin (n = 3, middle and right). Statistical significance was analyzed using one-way ANOVA followed by Tukey’s multiple comparisons test. \*\*\**p* < 0.001, \*\*\*\**p* < 0.0001. * vs age matched WT controls.

### Increased neuroinflammation in the hippocampus of *App* knock-in mice

Neuroinflammation has been previouly reported in the *App* knock-in mice ^12, 26^. In agreement, pathway analysis revealed activation of several key inflammatory signaling pathways including TNF, NF-κB, chemokine, Toll-like receptor signaling pathways, cytokine-cytokine receptor interaction and complement and coagulation cascade (**Figure 1E**). To identify affected cellular functions, Gene Ontology (GO) analysis was performed using significantly altered genes. One of the significantly altered GO biological process terms was inflammatory response (GO:0006954) (**Supplemental Table 1**). Analysis using various pathway enrichment tools as described in Material and Methods revealed a dramatic increase in DEGs involved in the neuroinflammatory response in the hippocampus of *App^NL-G-F^* mice from six months of age, whereas the inflammatory response was significantly increased in 18-month-old *App^NL-F^* mice (**Figure 2A**). To confirm the RNA-seq results, the expression of genes annotated to inflammatory response by GO term analysis (**Supplemental Table 1**) was measured by RT-qPCR. Upregulation of *Trem2*, *Clec7a, C4b and Ccl3* and downregulation of *Chrna7* were confirmed in both six and 12-month-old *App^NL-G-F^* mice, of which *Clec7a, C4b* and *Ccl3* expression continued to increase upon aging. *Trem2, C4b* and *Ccl3* were also increased in 18-month-old *App^NL-F^* mice (**Figure 2B**). These data support that the onset of neuroinflammation occurs earlier in *App^NL-G-F^* mice than in *App^NL-F^* mice, most likely due to the more aggressive Aβ pathology in the form of extracellular depositions of Aβ in *App^NL-G-F^* mice induced by the Arctic mutation. Inflammatory proteins including cytokines present in the brain can diffuse to CSF and may hence be utilized as potential markers for brain inflammation. To investigate if inflammatory proteins could be detected, and ultimately, if they are altered in CSF from late-stage *App* knock-in mice, CSF of 18-month-old mice were analyzed by Proximity Extension Assay using a Mouse Exploratory Panel. In agreement with the RNA-seq data showing an increased gene expression of *Ccl3*, Ccl3 protein levels are specifically increased in CSF of *App^NL-G-F^* mice. Other inflammatory markers Ccl5, Ccl20, Tnfrsf12a were altered differently in CSF of the two *App* knock-in mouse models, indicating differences in the inflammatory statuses in these two models. (**Figure 2C, Supplemental Table 2**).

**Figure 2.**
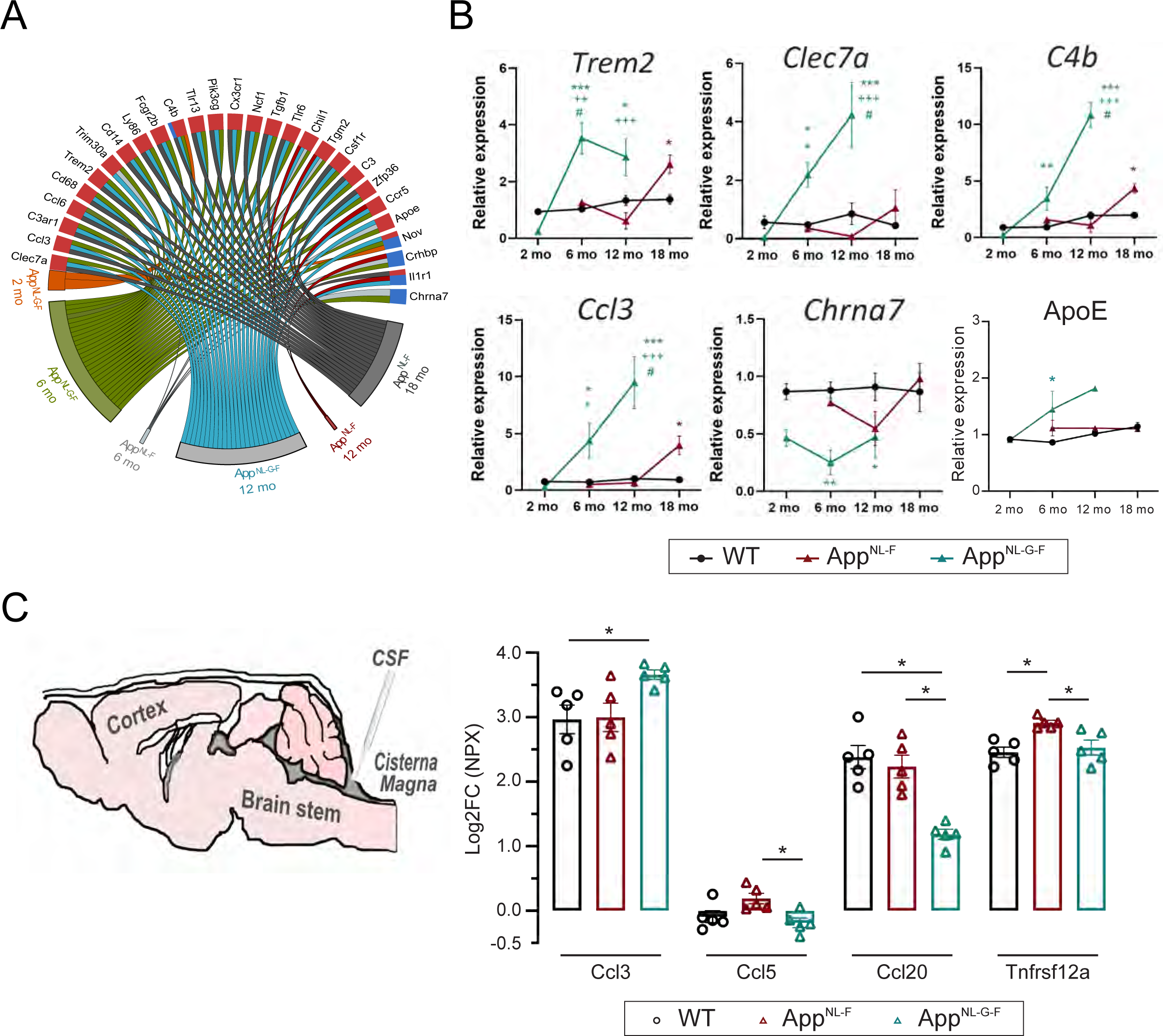
Neuroinflammation is a major pathologic phenotype in middle to late stage of *App* knock-in mice. (A) Chord plot of significantly altered genes (FDR < 0.1) related to inflammatory response. Color of the circle edge boxes indicate up-(red) or down-(blue) regulation. (B) Relative mRNA expression levels of selected inflammatory genes were normalized to *Tubb3* (n = 3). Statistical significance was analyzed using Kruskal-Wallis test followed by Dunn’s multiple comparison test. * vs age-matched WT mice, ^+^ vs age-matched *App^NL-F^* mice, ^#^ vs genotype-matched two-month-old mice, \**p* < 0.05, \*\**p* < 0.01, \*\*\**p* < 0.001. (C) CSF was withdrawn from cisterna magna through the dura mater of 18-month-old mice and analyzed by proximity extension assay technology using mouse exploratory panel to detect the changes in proteins related to inflammation. Statistical significance was analyzed using Kruskal-Wallis tests followed by Dunn’s multiple comparison test. \**p* < 0.05.

### Early symptomatic *App^NL-G-F^* mice display increased hippocampal OxPHOS activity along with calcium handling deficits

A large body of evidence has shown extensive mitochondria deficits in the brain of AD patients and in murine AD models; however, data available was mainly obtained at symptomatic stages when mitochondrial decline is evident and/or when APP is overexpressed ^27-29^. Notably, in this study, RNA-seq analysis and subsequent pathway analysis of the hippocampal transcriptomes revealed OxPHOS as one of the most upregulated pathways at early symptomatic stages in *App* knock-in mice (FDR= 0.0478 for *App^NL-F^* mice; FDR= 0.0085 for *App^NL-G-F^* mice) (**Figure 1E; Supplemental Figure S1**). A significant upregulation of nuclear-(*e.g., Ndufa1-3, Cox7c, Cox8b*) and mitochondrial DNA-(*e.g., mt-Nd4, mt-Atp6, mt-Atp8*) encoded genes was observed for four out of the five mitochondrial complexes in two-month-old *App^NL-G-F^* mice (**Fig. 3A-C, Supplemental Figure S2A**). *App^NL-F^* mice followed the same pattern of upregulation, but the number of metabolism-related DEG was remarkably lower (**Fig. 3C**). Upregulation of OxPHOS genes have been previously observed in hippocampus from mild cognitive impairment (MCI) patients, suggesting a compensatory-like mechanism at the early stages of disease ^30^. Both complex IV and ATP synthase are known to be targeted by Aβ ^31, 32^; concordantly, we observed that the most significantly altered genes with FDR<0.01 belong to complex V, followed by complex III and IV (**Figure 3B**), whereas RT-qPCR analysis showed a tendency of upregulation of some selected genes belonging to mitochondrial complexes I, III-V (**Supplemental Figure S2B**).

**Figure 3.**
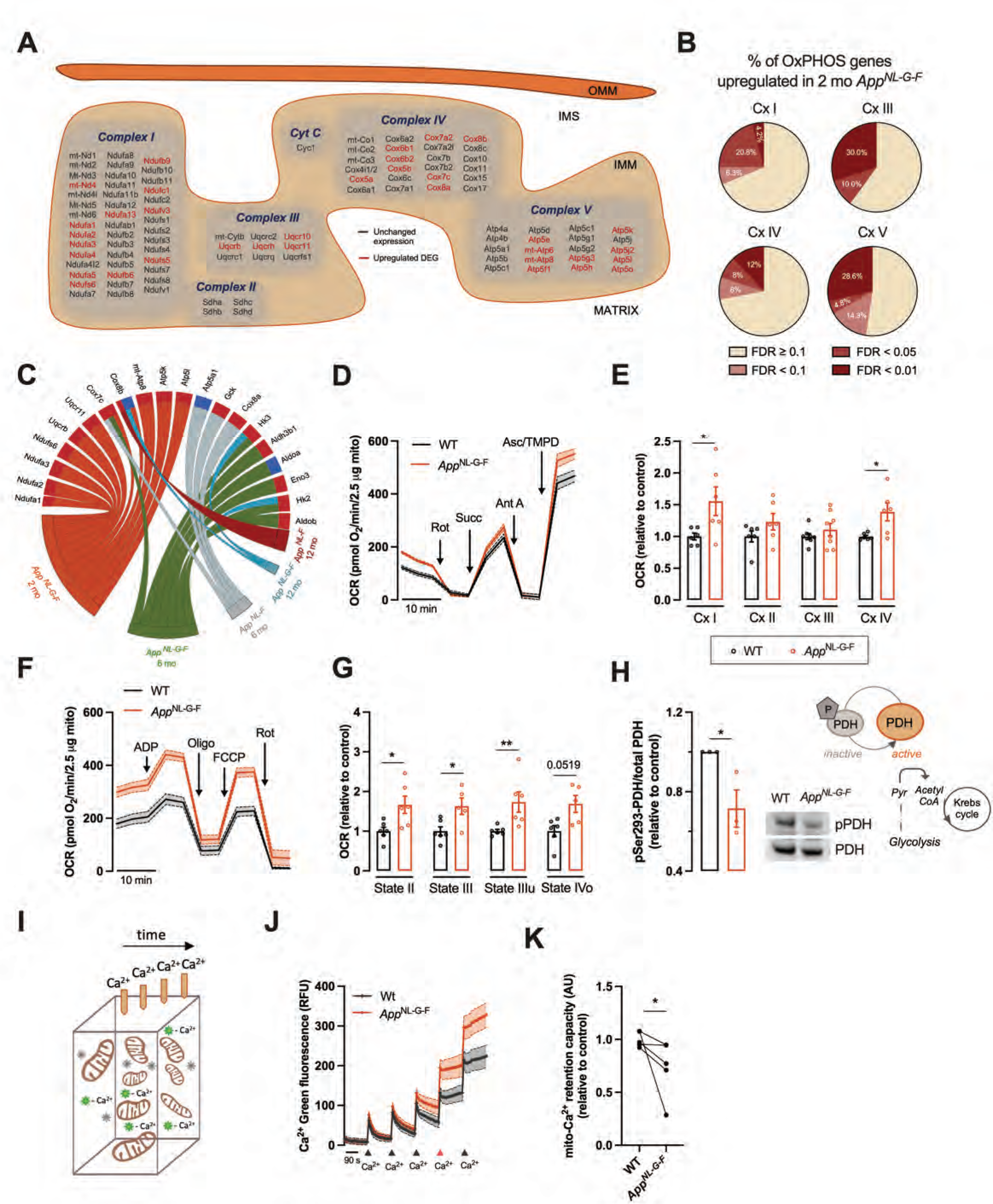
OxPHOS gene expression and activity were upregulated in early symptomatic *App^NL-G-F^* mice. (A) Organization of OxPHOS genes sorted by mitochondrial complex in the mitochondrial cristae. Significantly upregulated genes in two-month-old *App^NL-G-F^* mice are shown in dark red (FDR < 0.1). (B) Percentage of upregulated OxPHOS genes grouped by mitochondrial complex (Cx I, III, IV and V) in two-month-old *App^NL-G-F^* mice based on their FDR value. (C) Chord plot of significantly altered genes (FDR < 0.1) included in GO terms related to mitochondrial function. Color of the circle edge boxes indicate up-(red) or down-(blue) regulation. (D, E) Electron flow of uncoupled mitochondria isolated from two-month-old WT and *App*^NL-G-F^ mice was evaluated using the Seahorse apparatus. Mitochondrial complex inhibitors and substrates, 2 μM rotenone (Rot), 10 mM succinate (Succ), 4 μM antimycin A (Ant A), and 1 mM ascorbate (Asc)/100 mM TMPD, were sequentially injected to analyze the mitochondrial complex I-IV activities (*n* = 6 - 8). (F, G) OCR of mitochondria in coupled state, isolated from two-month-old WT and *App^NL-G-F^* mice using the Seahorse apparatus, to calculate the OCR of state II, state III induced by ADP (4 mM), state IIIu induced by FCCP (4 μM), and state IVo induced by oligomycin (3.2 μM, Oligo) (*n* = 5 - 6). (H) Phospho (p)-pyruvate dehydrogenase (PDH) (S293) and total PDH protein levels in hippocampal extracts were quantified by western blotting (*n* = 3). (I-K) Calcium uptake of hippocampal mitochondria was evaluated with the fluorescent probe Calcium-green. Five pulses of 10 μM CaCl_2_ were added to evaluate the mito-Ca^2+^ retention capacity (*n* = 3). Statistical significance was analyzed using non-parametric Mann Whitney test. \**p* < 0.05, \*\**p* < 0.01. OxPHOS: oxidative phosphorylation, OCR: oxygen consumption rate, IMM: inner mitochondrial membrane, IMS: inner mitochondrial space, OMM: outer mitochondrial membrane.

OxPHOS activity is tightly regulated by a series of events such as post-translational modifications and super-complexes formation. Therefore, it is highly contentious to speculate about mitochondria activity merely based on gene expression profile. To understand if the upregulation of TCA cycle and OxPHOS pathways could be translated into a functional upregulation, we isolated respiring-competent mitochondria from two-month-old *App^NL-G-F^* mice and evaluated the mitochondrial electron flow through electron transport chain (ETC) using the Seahorse XFe96 analyzer. Uncoupled mitochondria were incubated in malate and pyruvate to drive the activity of mitochondrial complex I and, subsequently, mitochondrial complexes activities were determined by sequential injection of mitochondrial inhibitors and substrates (**Figure 3D**). We observed an increase of 55% in complex I (*p* = 0.0260) and 39% in complex IV activities (*p* = 0.0411), with no changes in complex II and III from two-month-old *App^NL-G-F^* hippocampal mitochondria (**Figure 3E**). Complex I and IV activities highly correlated with RNA-seq data, in which 31.3% and 28% of genes analyzed from complex I and IV, respectively, were upregulated (**Figure 3B**). We further evaluated mitochondrial respiration to assess if elevated OxPHOS drive ATP production through complex V (**Figure 3F, G**). Increased state II mitochondria respiration (*p* = 0.0238) (**Figure 3G**), together with boosted pyruvate dehydrogenase activation (*p* = 0.0242) (**Figure 3H**), a key enzyme that links glycolysis to TCA cycle in which inhibitory phosphorylation is associated with a decay in activity, confirms an overall upregulation of mitochondrial function in the hippocampus of two-month-old *App^NL-G-F^* mice. Addition of ADP to fuel complex V further led to an increase in state III (*p* = 0.0173), suggesting an increased capacity to generate ATP (**Figure 3G**), possibly related to increased levels of ETC proteins. Interestingly, augmented state IV respiration induced by oligomycin (IVo) was observed in *App^NL-G-F^* mitochondria (*p* = 0.0519) (**Figure 3G**), suggesting an increase in leakage of electrons from the ETC. These data support the hypothesis that enhanced respiration in two-month-old *App^NL-G-F^* hippocampus can potentiate oxidative stress. Concordantly, GO clustering for biological processes in the two-month-old *App^NL-G-F^* mice revealed upregulation of mitochondrial respiratory complex I assembly, together with changes in oxidation-regulation processes (**Supplemental Figure S2C**). These data suggests that enhanced respiration in the hippocampus of two-month-old *App^NL-G-F^* mice can potentiate reactive oxygen species (ROS) production and consequent oxidative imbalance. Since the rate of ATP synthesis dynamically matches with mitochondrial calcium (Ca^2+^) homeostasis, we performed a mitochondrial calcium (Ca^2+^) handling assay to understand if increased electron flow through the mitochondrial ETC correlates with enhanced function. Isolated mitochondria from *App^NL-G-F^* hippocampi were incubated with a visible light-excitable Ca^2+^ indicator, and several Ca^2+^ pulses were consecutively performed to test the mitochondrial Ca^2+^ handling capacity (**Figure 3I, J**). Cumulative fluorescence intensity in the experimental media indicates that *App^NL-G-F^* mitochondria lose their capacity to accumulate Ca^2+^ much faster than WT mitochondria (**Figure 3J**). Further analysis of mitochondrial Ca^2+^ retention capacity confirmed previous results (*p* = 0.0317) (**Figure 3K**). Overall, these data indicate that mitochondria isolated from hippocampus of two-months-old *App^NL-G-F^* mice display increased OxPHOS, potentially as compensatory response, which may contribute to unbalanced oxidative status and maladaptive Ca^2+^ handling at early stages.

### Mitochondrial deficits and synaptic structural disorganization characterize late symptomatic *App^NL-G-F^* mice

Based on our transcriptomic data and the subsequent pathway analysis, the mitochondrial metabolism in the hippocampus of *App^NL-G-F^* mice started to decay before six months of age and the decline continued till further aging (FDR= 0.041 at six mo; FDR= 0.006 at 12 mo for OxPHOS) (**Figure 1E**). ATP hydrolysis and axonal mitochondrial transport were among the ten most downregulated processes, as identified by GO analysis, in the hippocampus of six-month-old *App^NL-G-F^* mice, confirming that a shift in energy metabolism occurs between two to six months of age (**Figure 4A**). Moreover, GO analysis of six and 12-month-old *App^NL-G-F^* mice also revealed deficits in synapse organization which is associated with downregulated synaptic vesicles transport and exocytosis (**Figure 4A**). To functionally validate the decay in energy metabolism, we evaluated mitochondrial oxygen consumption rate (OCR) of mitochondria from hippocampus of 12-month-old *App^NL-G-F^* and WT mice (**Figure 4B-E**). Consistent with GO analysis, we observed a severe impairment of mitochondrial complexes I, II and III, as shown by reduced activities being 49% (*p* = 0.0079), 35% (*p* = 0.0317) and 34% (*p* = 0.0317), respectively, for each complex, whereas no significant impairment of mitochondrial complex IV activity was detected **(Figure 4C)**. Surprisingly, declined activities of mitochondrial complexes did not translate into decreased state II or state III respiration (**Figure 4D, E**), suggesting that mitochondrial respiration is tightly regulated to sustain ATP production. Nevertheless, mitochondria in the hippocampus of *App^NL-G-F^* mice exhibited deficient respiration when they were forced to respire at maximal capacity upon depolarization (state IIIu) (*p* = 0.0476) (**Figure 4D, E**). In addition, analysis of electron micrographs of hippocampus from 10- to 12-month-old *App^NL-G-F^* mice further demonstrated a reduction of mitochondrial profile number at the pre-synaptic terminals (*p* = 0.0227) (**Figure 4F, G, Supplemental Figure S3**). Analysis of the endoplasmic reticulum (ER) network also revealed an increase in the ER aspect ratio, a measure of elongation, together with decreased thickness in *App^NL-G-F^* CA hippocampus (*p* < 0.0001) (**Figure 4H, Supplemental Figure S3**), in agreement with the GO analysis indicating alterations in ER tubular network organization at this age (**Figure 4A**). The extended ER structure can partially explain why the *App* knock-in mice display increased mitochondria-ER contacts sites (MERCS) per mitochondria and consequent mitochondrial fragmentation ^33^. Furthermore, EM analysis illustrated abnormally enlarged pre-synaptic areas (*p* < 0.0366) (**Figure 4F, I**), increased number of synaptic vesicles (*p* < 0.0001) (**Figure 4F, J**), decreased post-synaptic density thickness (*p* < 0.0001) (**Figure 4F, K**). Around Aβ plaques, the increase in presynaptic area and the number of synaptic vesicles is further pronounced and in this area an accumulation of double membrane autophagic vacuoles (AVs) occurred (**Figure 4F, Supplemental Figure S3**). Abnormally enlarged pre-synaptic areas (*p* = 0.0070) and decreased post-synaptic density thickness (p < 0.0001), but not a reduction of mitochondrial profile number, were similarly observed in hippocampus from 22- to 24-month-old *App^NL-F^* mice. An accumulation AVs was additionally observed around Aβ plaques in hippocampus of *App^NL-F^* mice as we previously reported ^34^ (**Supplemental Figure S3**).

**Figure 4.**
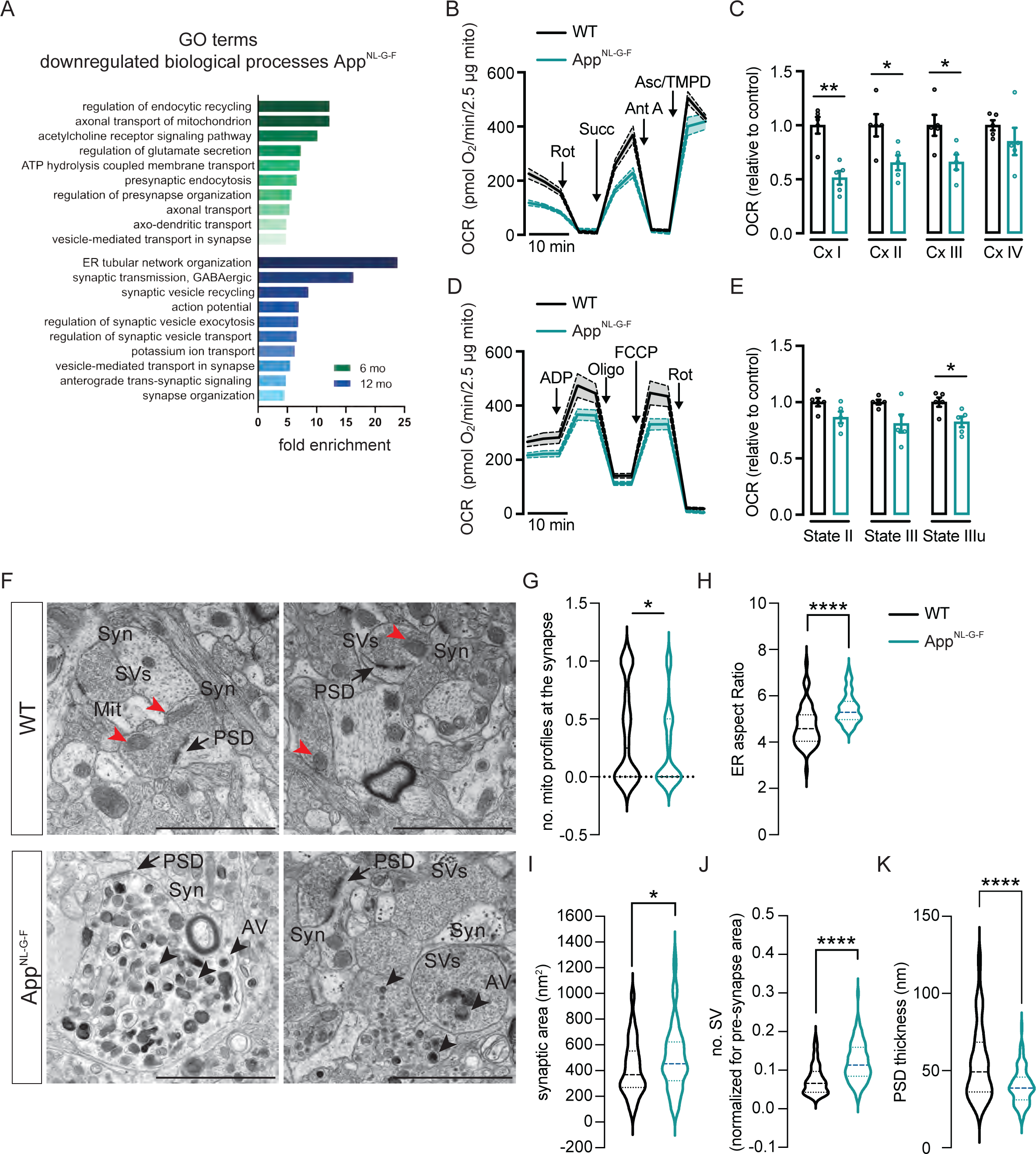
Decreased mitochondrial respiratory output in synapsis is paralleled by morphological alteration of ER, autophagosome accumulation and aberrant synapse assembly. (A) Significantly downregulated GO biological process terms associated with synaptic and transport functions in *App^NL-G-F^* mice. (B, C) Electron flow of uncoupled mitochondria and (D, E) OCR of coupled mitochondria isolated from 12-month-old WT and *App*^NL-G-F^ mice (n = 5). (F) Electron microscopy images of hippocampal CA1 from 10- to 12-month-old WT and *App^NL-G-F^* mice (n = 4, with an average of 50 cells and 70 synapses analyzed per genotype). Black arrowhead: AV, red arrowhead: Mitochondria, black arrow: postsynaptic density. Scale bar: 2 µm. Quantification of mitochondrial profiles in the synapses (G), ER aspect ratio (H), synaptic area (I), number of synaptic vesicles normalized by pre-synaptic area (J) and post-synaptic thickness (K) in WT and *App^NL-G-F^* mice. Statistical significance was analyzed using non-parametric Mann Whitney test. \**p* < 0.05, \*\**p* < 0.01, \*\*\*\**p* < 0.0001. AV: autophagic vesicle, Mit: Mitochondria, PSD: postsynaptic density, SVs: synaptic vesicles, Syn: synapse.

### Autophagic and synaptic vesicles accumulate at the presynapse around Aβ plaques in aged *App knock-in* mice

Having found a massive accumulation of Avs in 12 mo NLF and 24 NLF mice prompted us to further investigate the autophagy statues in the mice, Macroautophagy, here referred to as autophagy, is an intracellular self-digesting system which is impaired in most neurodegenerative disorders including AD, and directly involved in the Aβ metabolism ^35, 36, 37^. We further assessed the transcriptional alterations of autophagy genes in *App* knock-in mice. Consistently, the mTOR signaling pathway, which is a key negative regulator of autophagy and hyperactivated in AD brains ^38^, was activated both in 12-month-old *App^NL-F^* and *App^NL-G-F^* mice (FDR= 0.0167 for *App^NL-F^*; FDR= 0.0121 for *App^NL-G-F^*) (**Figure 1E**). In addition, the regulation of autophagy pathway was slightly downregulated in 12-month-old *App^NL-G-F^* mice (FDR = 0.0895) (**Figure 1E**). We also identified significantly altered autophagy genes already at six months of age including *Trim30a, Rubcnl, Vamp8, Lamp2* and *Rab7b* annotated by GO terms and confirmed by RT-qPCR (**Figure 5A, B**). Furthermore, the RNA-seq data revealed a strong upregulation of the GTPase Rab7b, which binds to and negatively regulates autophagy through acting on Atg4B thereby controlling LC3I lipidation, size of autophagosomes and ultimately autophagy flux^66^.

**Figure 5.**
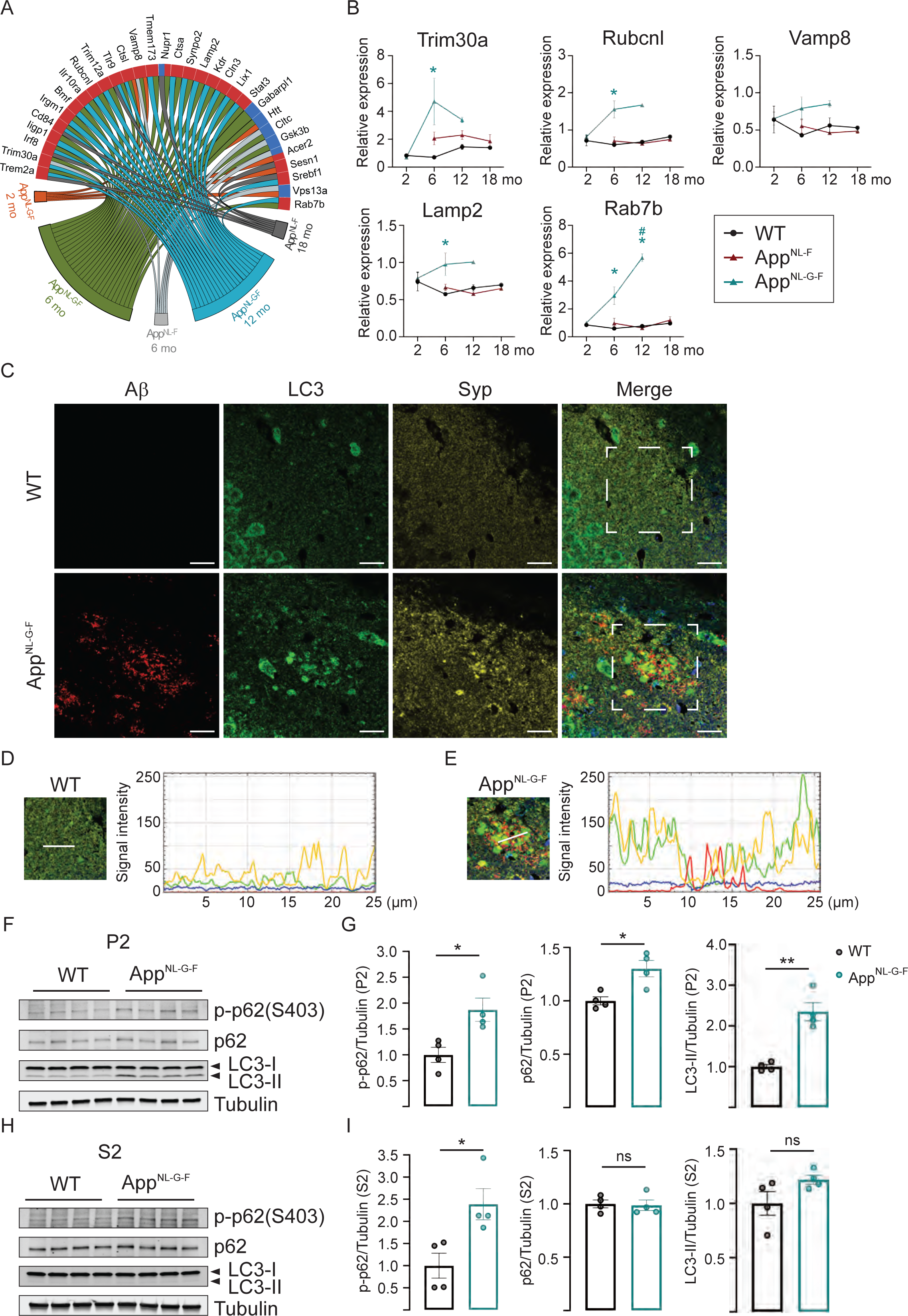
Impaired synaptosomal autophagy in aged *App^NL-G-F^* mice is observed especially around Aβ plaques. (A) Chord plot of significantly (FDR < 0.1) DEGs related to autophagy. Color of the circle edge boxes indicate up-(red) or down-(blue) regulation. (B) The relative mRNA expression of selected genes was normalized to *Tubb3* (n = 3). Statistical significance was analyzed using Kruskal-Wallis tests followed by Dunn’s multiple comparison test. \**p* < 0.05. * vs age matched WT mice, ^#^ vs 2-month-old genotype-matched mice. (C) Co-immunofluorescence staining of Aβ, LC3 and synaptophysin with 12 months-old WT and *App^NL-G-F^* mice (n = 4). Red: Aβ, Green: LC3, Yellow: Synaptophysin, Blue: nucleus. Scale bar: 20 µm. (D, E) Signal intensity of Aβ (Red), LC3 (Green), Synaptophysin (Yellow) and nucleus (Blue) under the white line of WT and *App^NL-G-F^* brain staining gated in C. (F-I) Phospho-p62 (S403), total p62, LC3-I and LC3-II protein levels in hippocampal crude synaptosomal fraction (P2) or soluble fraction (S2) were visualized by Western blotting (n = 4). Protein levels were normalized to β3-tubulin. Statistical significance was analyzed using unpaired t test. \**p* < 0.05, \*\**p* < 0.01.

Taking all the data together including RNA-seq and EM analysis, this indicates that autophagy is altered especially in the *App^NL-G-F^* mice at an age when a robust Aβ pathology is established, and we therefore continued to investigate autophagy in 12-month-old *App^NL-G-F^* mice. To further analyze the autophagy alteration and its relation to Aβ plaque pathology and synaptic alterations, we assessed autophagy by triple immunofluorescence staining against microtubule-associated protein 1 light chain 3 (MAP1LC3), which is widely used as a maker for phagophores and autophagosomes, Aβ and the synaptic marker synaptophysin in 12-month-old *App^NL-G-F^* and WT mice. Granular staining of LC3 was observed surrounding the Aβ plaques, possibly corresponding to the accumulation of Avs in the dystrophic neurites around the Aβ plaques (**Figure 5C**). Interestingly, an accumulation of synaptophysin was found around the Aβ plaques which partially colocalized with the granular LC3 accumulation (**Figure 6D, E**), supporting the EM data showing enlarged presynaptic terminal containing Avs (**Figure 4F, Supplemental Figure S3**). Next, we evaluated LC3 and p62/SQSTM1 levels in crude synaptosomal fraction which contained mainly presynaptic terminal and postsynaptic membrane as validated by synaptophysin and postsynaptic marker postsynaptic density protein 95 (PSD95). The ratio of synaptophysin to PSD95 was significantly increased in *App^NL-G-F^* brain caused by a reduction of PSD95, as previously reported ^39^ (**Supplemental Figure S4A-C**). LC3 exists in cytosol as a soluble form, LC3-I, and is converted to LC3-II by conjugation with phosphatidylethanolamine upon insertion into the autophagosomal membrane. Hence LC3-II is used as the maker for phagophores and autophagosomes. P62 is a receptor for facilitating autophagic degradation of ubiquitinated substrates ^40^. Notably, LC3-II and p62 were both significantly increased in the hippocampal synaptosomal fraction of *App^NL-G-F^* mice whereas no significant differences of LC3-II and p62 levels were detected in the soluble fraction (**Figure 5F-I**). In agreement with the autophagy alteration occurring from six-months-of-age in the *App^NL-G-F^* mice as determined by the transcriptomic analysis, no autophagy alteration was observed in two-month-old *App^NL-G-F^* mouse brain as determined by p62 and LC3 western blot analysis of synaptosomal fractions and LC3 immunostaining (**Supplemental Figure S4D-H**). Phosphorylation of p62 at serine 403 (S403) stabilizes the binding between p62 and ubiquitinated protein, therefore enhances degradation of ubiquitinated proteins by autophagy ^41^. As expected, p-p62 (S403) levels were increased in crude synaptosomal fraction of *App^NL-G-F^* mouse brains (**Figure 5F, G**). The number of autophagosomes can be increased either by the activation of autophagosome production or by the inhibition of autophagosome degradation. To reveal whether autophagy activation is induced in the synapses in the *App^NL-G-F^* mice, we investigated two phosphorylation sites, S757 and S555, of ULK1, the main regulator of autophagy initiation ^42 43^. Phosphorylation at S757, which inhibits ULK1, was significantly reduced in the synaptosomal fraction of *App^NL-G-F^* mice (**Supplemental Figure S5A-E**). However, the level of total ULK1 was also decreased in *App^NL-G-F^* mice and we conclude that no change in synaptosomal autophagy initiation is present in the 12-month-old *App^NL-G-F^* mice. Instead, an inhibition of synaptosomal Avs degradation around the Aβ plaques may explain the accumulation of Avs in aged *App^NL-G-F^* mice which is supported by a decrease in gene expression of several lysosomal vATPases (**Supplemental Figure S5A-E, Supplemental Table 3**)

## DISCUSSION

In this study we investigated the effects of Alzheimer-associated Aβ amyloidosis by analyzing the transcriptome of hippocampus, which is a key region for memory formation, of two *App* knock-in mouse models exhibiting different degrees of AD-like phenotypes including Aβ pathology, neuroinflammation and cognitive impairments. The analysis of the 46,000 obtained transcripts revealed that both *App^NL-F^* and *App^NL-G-F^* mice exhibited substantial alterations in their transcription profiles upon progression of the pathologies and especially pronounced in *App^NL-G-F^* mice which exhibited more than 2000 DEGs. Analysis of the affected pathways revealed that several key cellular functions were significantly affected. These included 1) early changes in mitochondrial function, 2) onset of neuroinflammation, 3) autophagy impairment. These changes ultimately lead to an abnormal pre-synaptic organization likely contributing to the synaptic impairment and memory decline in the *App* knock-in mice. By combing transcriptome data with functional analysis, we showed that one of the first events observed in early phase of the development of the AD-associated pathologies is a hypermetabolism mainly characterized by upregulation of OxPHOS. Previously we showed that Aβ can be internalized into mitochondria ^44^ and *ex vitro* studies indicated that both the extracellular domain of APP and Aβ can interact with ATP synthase subunit α and partially inhibit extracellular ATP generation ^45^. The upregulation of mitochondrial ETC gene expression resulting in an activity increase preced a strong Aβ amyloidosis in *App* knock-in mice may occur as a compensatory mechanism to bypass deficits in ATP generation, as previously suggested ^46^. A significant upregulation in OxPHOS genes in MCI subjects compared to control or AD cases has been described before, with genes belonging to complexes I, III, IV and V being the most affected, but not those in complex II ^46^. Additionally, data from two-month-old APP Tg2576 mice also showed that most of the upregulated DEG were related to mitochondrial energy metabolism and apoptosis. Consistently, mitochondria from two-month-old *App^NL-G-F^* mice exhibited higher susceptibility to Ca^2+^ overload, which has been reported to be associated with neuronal death and cognitive decline in different AD models ^50, 51^. Several underlying mechanisms are probable causes ^50^, including increased Ca^2+^ shuttling from ER to mitochondria due to upregulated MERCS, which we have reported before in *App* knock-in mice ^33^. Increased MERCS can account for both Ca^2+^ overload susceptibility and enhanced OxPHOS activity dependent on TCA cycle. We find that OxPHOS upregulation is one of the most primary characteristics of neuronal vulnerability leading to manifested oxidative damage ^52^, further confirmed by increased H^+^ leakage across the mitochondrial inner membrane and upregulation of oxidation-reducing processes in our RNA-seq data. Interestingly, both mitochondrial ETC and sustained mitochondrial-derived ROS generation are necessary for inflammasome activation. Neuroinflammation in *App* knock-in mice have previously been described to comprise both astrocytosis and microgliosis further confirmed here by increases in AD-associated *ApoE* and *Trem2* and by activated neuroinflammatory pathways including TNF signaling, toll-like receptor signaling, cytokine signaling and activation of the complement system ^12, 53, 26, 54, 55^. Interestingly, a large-scale proteomic analysis of AD brains revealed that increased sugar metabolism in astrocytes and microglia emerged as one of the network modules most significantly associated with AD pathology and cognitive impairment ^28^. Therefore, it is plausible that a feedback cycle exists between ECT activity and inflammatory cascades. Although the neuroinflammatory response is manifested upon aggravation of the Aβ pathologies in *App* knock-in mice, starting after two months-of-age in *App^NL-G-F^* mice and 12-months-of age in the *App^NL-F^* mice, a recent study has shown increased pro-inflammatory cytokines such as TNF-α, IL-6 and IL17A/F as early as two months-of-age in the brains of *App^NL-F-G^* mice, in agreement with an early microglial activation. Upon aging, the levels of anti-inflammatory molecules increase, however, inflammation remains unresolved in the late stage of pathology. The gene expression of CCL3, a key component of the complement system, was increased from six-month-of age in *App^NL-G-F^* mice and from 18-month-of-age in *App^NL-F^* mice. Interestingly, this translation of increased CCL3 gene expression to the CSF suggests its potential as CSF biomarker to trace neuroinflammation in the brain. In contrast to the increase in OCR detected in young mice, old *App^NL-G-F^* mice revealed a severe diminution of mitochondrial function, although ATP production was spared likely as a result of the assembly of respiratory super complexes ^57^. Considering that GO and pathway analysis of the transcriptomes additionally pointed towards a synaptic impairment prompted a thorough analysis of the synapse. Indeed, morphological characterization by EM confirmed a number of alterations (**Figure 4F-K**). Firstly, a decrease of synaptic mitochondria was noted. This was paralleled by enlarged presynaptic areas with a drastically increased number of synaptic vesicles. This abnormal enlargement of the synaptic vesicle pool could be due to Aβ-induced disruption of vesicle fusion, altered turnover of vesicles or impaired vesicle-mediated transport, as indicated by GO analysis (**Figure 4A**). In addition, decreased mitochondrial-derived local ATP supply could also contribute to the accumulation of synaptic vesicles since the synaptic vesicle cycle is a major consumer of ATP ^58^. Secondly, the reduction of post-synaptic density thickness and PSD95 protein level in the synapses of *App^NL-G-F^* mice suggests an impairment of synaptic vesicle exocytosis, resulting in a large inactive pool of synaptic vesicles, since PSD morphology is linked presynaptic terminal function ^59^ (**Figure 4F, K, Supplemental Figure S3**). Even though the *App^NL-F^* mice exhibit less pronounced Aβ pathology, similar synaptic alterations were observed including increased presynaptic regions and decreased PSD. However, no changes in synaptic mitochondria were found which correlates with less pronounced transcriptomic changes in mitochondrial related genes and pathways. These substantial morphological alterations in the synapse, which are ATP-dependent dynamic structures, most likely reflect an impaired synapse and neuronal transmission leading to the memory impairment in the *App* knock-in mice. Indeed, mushroom spine loss has been previously reported in the *App* knock-in mice ^60^ and electrophysiological characterization of *App^NL-G-F^* mice have shown a desynchronization and lowered amplitude of kainite-induced gamma oscillation ^61^. It is also noteworthy that activated microglia has a negative impact on synapses in AD ^62^.

Interestingly, an accumulation of AVs was observed in the direct vicinity of the synaptic vesicles. This implies a role of autophagy in the maintenance of protein homeostasis in the synapses and which is further supported by RNA-seq data, confirming a dysfunctional autophagy in aged *App^NL-G-F^* mice. The massive accumulation of AVs around Aβ plaques in dystrophic neurites and synapses is in line with an accumulation of AVs in AD brain^37^. Autophagosomes are formed in the presynaptic region as well as in the cell soma and the dendrites and transported retrogradely to the cell body for degradation through fusion to lysosomes ^63^. Notably, retrograde transport of autophagosomes is disturbed in AD ^63^ and in the *App* knock-in mice including axonal transport as identified by the pathway analysis. Alterations in transport could also be due to the reduced ATP levels caused by impaired mitochondrial function, which in turn could be due to lowered mitophagy causing a vicious cycle, but further functional studies are required to reveal the underlying mechanisms of autophagosome accumulation in *App^NL-G-F^* mice ^64^. The transport machinery ultimately delivers its content to lysosomes where an acidification is required for the final degradative step. This is largely controlled by vATPases, the impairment of which has previously been linked to FAD-causing PS1 mutations and altered early in the development of Aβ pathology ^65^. Our data revealed a clear autophagy inhibition associated with an Aβ plaque pathology and in the synapses which was paralleled by a decreased gene expression of a several vATPases were observed in six-month-old *App^NL-G-F^* mice indicating lysosomal alterations, but its cellular origin of this decrease remains to be established. The presynaptic autophagy-impairment potentially contributes to the enlargement of pre-synaptic area and massively packed synaptic vesicles. Previous data from pulse chase experiments in *App* knock-in mice indeed revealed an impaired protein metabolism of presynaptic proteins ^66^. Newly synthesized synaptic vesicle proteins are involved in neurotransmitter secretion and aged synaptic vesicle proteins are found in synaptic vesicles in the inactive reserve pool ^67^. Impaired autophagy may influence the balance between the active recycling and the inactive reserve pool, for example through decreased synaptic vesicle degradation, and subsequent synaptic vesicle accumulation. This hypothesis is supported by evidence showing that synaptic vesicle number and synaptic vesicle protein turnover are regulated by autophagy ^68^. Taken together, we have deciphered the temporal appearance of some of the AD-associated pathological alterations using the *App* knock-in mouse models which includes a striking early compensatory mitochondrial hyperactivity followed by a strong neuroinflammation and autophagic decline leading to faulty synapses. Thus, we propose that early interventions aiming at strengthening the synapse may serve as potential therapeutic strategies for AD through 1) mitochondrial improvement 2) damping neuroinflammation 3) improving protein homeostasis via autophagy.

## ACKNOWLEDGEMENT

We thank Takaomi Saido and Takashi Saito at RIKEN Center for Brain Science for providing *App* knock-in mice. We thank the EM facility in Huddinge Hospital (EMil) and Maho Hamasaki and Tamotsu Yoshimori at Osaka University for valuable help with EM and the Beta Cell *in-vivo* Imaging/Extracellular Flux Analysis core facility, supported by the SRP Diabetes, and Dr. Noah Moruzzi for the help with Seahorse experiments. We are grateful for financial support from: Hållsten Research Foundation (PN), Swedish Research Council (PN, MA), Swedish Brain Foundation (PN, MA), Torsten Söderberg Foundation (PN), Sonja Leikrans donation (PN), The Erling-Persson Family Foundation (PN) and the Swedish Alzheimer Foundation (PN, MA, MS, LN). China Scholarship Council (RJ), Gun and Bertil Stohne’s Research Scholarship (RJ). LN and MS are funded by the Strategic Research Program in Neuroscience (StratNeuro) funding for postdoctoral researchers. TOYOBO biotechnology foundation (MS). Japanese Society for the Promotion of Science (NSL).

## Data availability

The RNA-seq data will be deposited to the NCBI Sequence Read Archive (SRA).

## Author contributions

LN, MS and E. Bereczki performed RT-qPCR validations; LN designed, performed and analyzed mitochondrial experiments; MS and NSL performed mitochondrial experiments; MS designed experiments related to autophagy; MS, E. Berger and VLF performed and analyzed, IHC and biochemical experiments related to autophagy and synapsis; MS and NSL prepared samples for electron microscopy and acquired micrographs which MS, LN and GD analyzed. RJ collected and analyzed CSF; PN and MA performed animal dissections and collection of tissues; JL performed RNA sequencing; XL and E. Bereczki did bioinformatic analysis. LN, MS, CMP, MA and PN were responsible for the coordination and experimental design. MA and PN secured funding. LN, MS, E. Bereczki, MA, PN wrote the manuscript with the agreement of all the other authors.

---

**Supplemental Figure 1. Transcriptome pathway analysis.** Pathway analysis was performed using Gene Set Enrichment Analysis (GSEA) method based on the KEGG dataset. The *p*-value of each pathway was converted to Z score. Significantly up- and downregulated pathways after enrichment have an absolute value of Z scores ≥ 1.96. The pathways are arranged in hierarchical order. Genes highlighted in red is upregulated and blue is downregulated.

**Supplemental Figure 2. Mitochondrial-related heatmap, GO clustering and gene expression analysis reveals significant changes already at two months in *App^NL-G-F^* mice.** (A) Heatmap showing gene expression converted to Z score from the KEGG pathway dataset ‘Oxidative Phosphorylatio’. Genes belonging to mitochondrial complexes (I-V) are arranged together. Genes highlighted in red are up- and blue are downregulated. (B) Relative mRNA expression of selected genes belonging to different mitochondrial complexes (Cx). Data was normalized to *Tubb3* (n = 3). (C) Gene Ontology clustering plot for biological processes. The color of the circles corresponds to log_10_(p-value) changes. Statistical significance was analyzed using Kruskal-Wallis tests followed by Dunn’s multiple comparison test. \**p* < 0.05 vs age matched WT controls, ^#^ vs two-month-old *App^NL-G-F^* mice.

**Supplemental Figure 3. Overview of altered synaptic morphology in *App^NL-G-F^* mice.** (A) Merged electron microscopy images of hippocampal CA1 from 12-month-old WT and *App^NL-G-F^* mice. Scale bar represents 2 µm in merged images, and 1 µm in zoom-in images. (B) Quantification of ER thickness of ten- to 12-month-old WT and *App^NL-G-F^* mice (n = 4, with an average of 50 cells and 70 synapses analyzed per genotype). (C) Merged electron microscopy images of hippocampal CA1 from 22- to 24-month-old WT and *App^NL-F^* mice. Scale bar represents 2 µm in merged images, and 1 µm in zoom-in images. Quantification of mitochondrial profiles in the synapses (D), synaptic area (E), number of synaptic vesicles normalized by pre-synaptic area (F) and post-synaptic thickness (G) in WT and *App^NL-F^* mice (n = 3 including one WT male, with an average of 50 cells and 70 synapses analyzed per genotype). Statistical significance was analyzed using non-parametric Mann Whitney test. \*\**p* < 0.01, \*\*\*\**p* < 0.0001. AV: autophagic vesicle, Mit: Mitochondria, PSD: postsynaptic density, SVs: synaptic vesicles, Syn: synapse.

**Supplemental Figure 4. Unaltered synaptosomal autophagy in two months old *App^NL-G-F^* mice.** (A, B) Protein levels of synaptophysin and PSD95 in hippocampal crude synaptosomal fraction (P2) or soluble fraction (S2) from 12-month-old WT and *App^NL-G-F^* mice were detected by western blotting (n = 4). (C) Synaptophysin and PSD95 protein level, and synaptophysin vs PSD95 ratio in P2 fraction. Statistical significance was analyzed using unpaired *t*-test. \*\**p* < 0.01. (D) Co-immunofluorescence staining of Aβ and LC3 with two-month-old WT and *App^NL-G-F^* mice (n = 4). Red: Aβ, Green: LC3, Blue: nucleus. Scale bar: 100 µm. (E-H) Total p62, LC3-I and LC3-II protein levels in P2 or S2 fraction from two-month-old WT and *App^NL-G-F^* mice were quantified by western blotting (n = 4). Protein levels were normalized to β3-tubulin. Statistical significance was analyzed using unpaired *t*-test.

**Supplemental Figure 5. No change in synaptosomal autophagy initiation in 12-month-old *App^NL-G-F^* mice. (A)** Phosphorylated ULK1 (S757) protein levels and (B) phosphorylated ULK1 (S555) and total ULK1 protein levels as well as their ratios in hippocampal crude synaptosomal fraction (P2) from 12-month-old WT and *App^NL-G-F^* mice were detected by western blot (n = 4). Statistical significance was analyzed using unpaired *t*-test. \**p* < 0.05, \*\**p* < 0.01.

