## Supplementary material for "Early mitochondrial dysfunction proceeds neuroinflammation, synaptic alteration, and autophagy impairment in hippocampus of *App* knock-in Alzheimer mouse models": Supp Fig 1-5

### Supplemental Figure 1

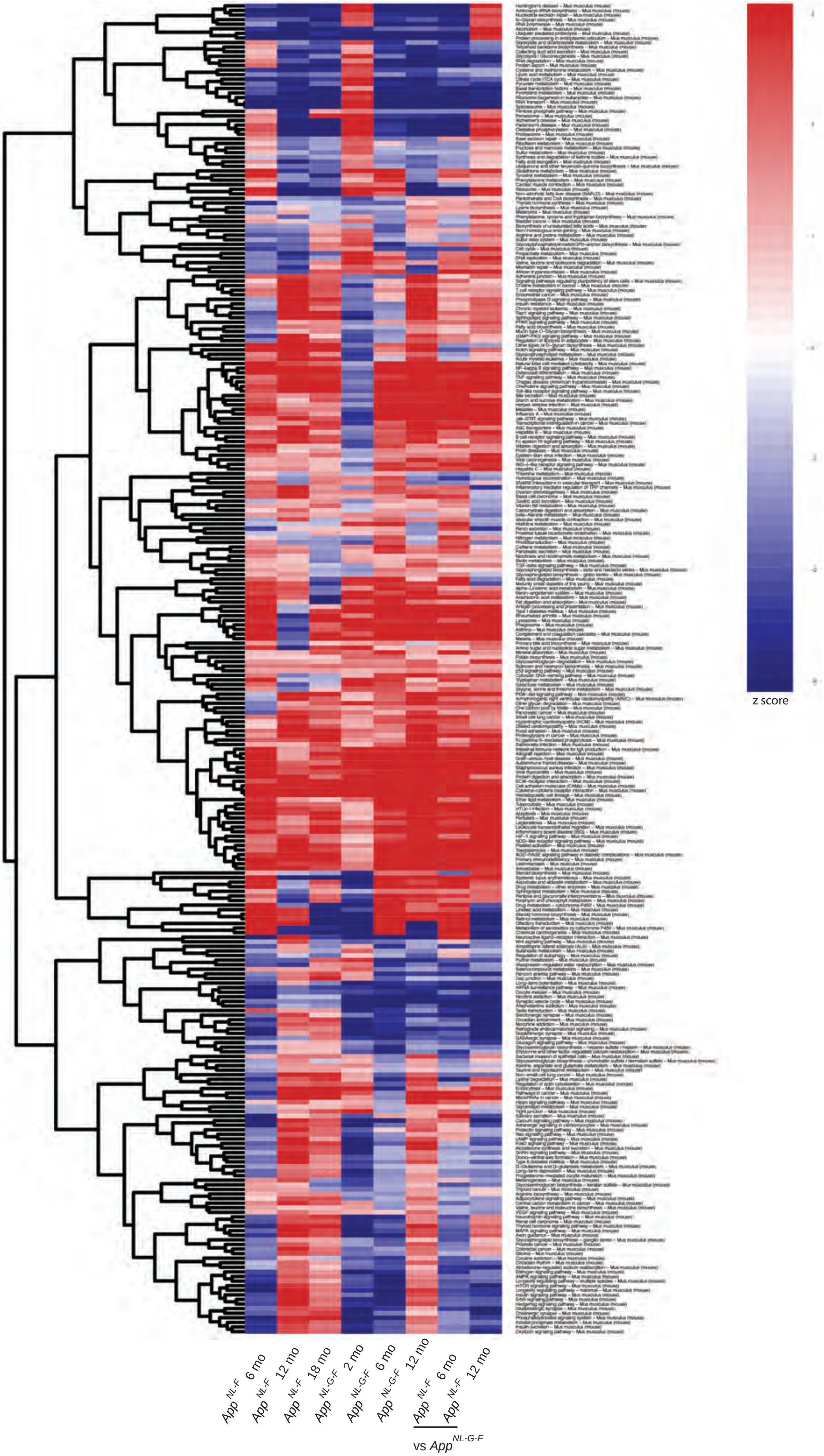

#### Supplemental Figure 2

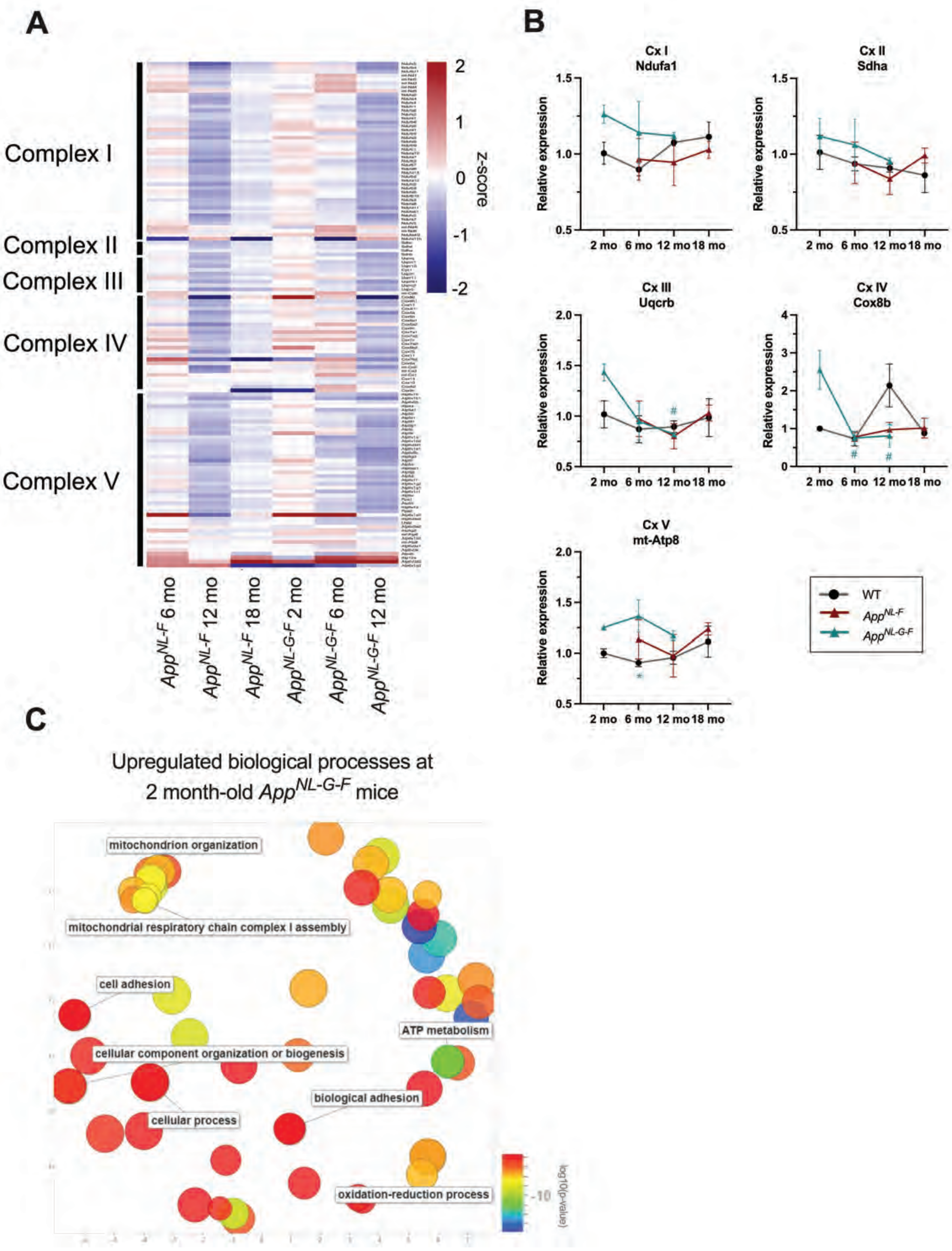

### Supplemental Figure 3

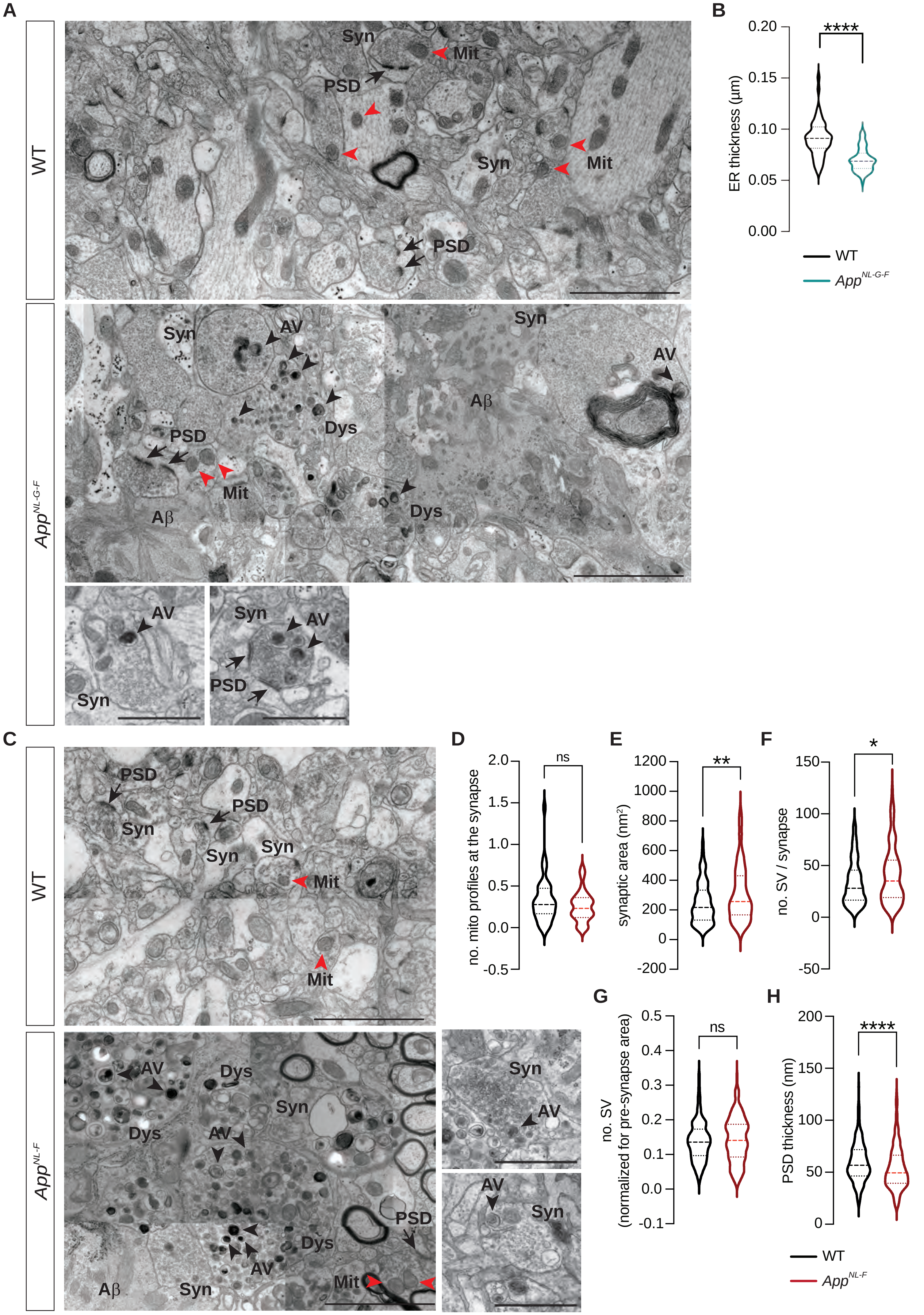

### Supplemental Figure 4

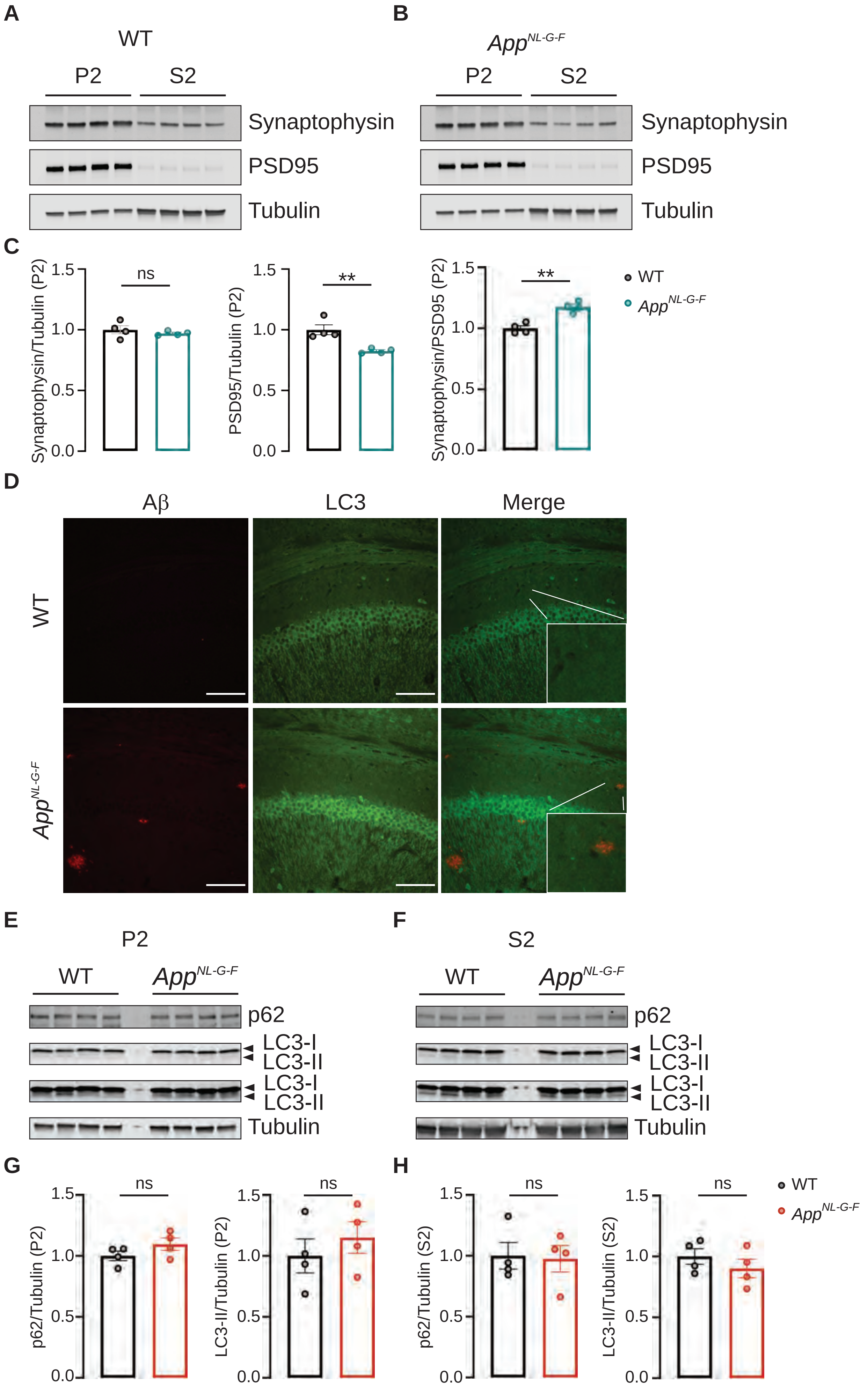

### Supplemental Figure 5

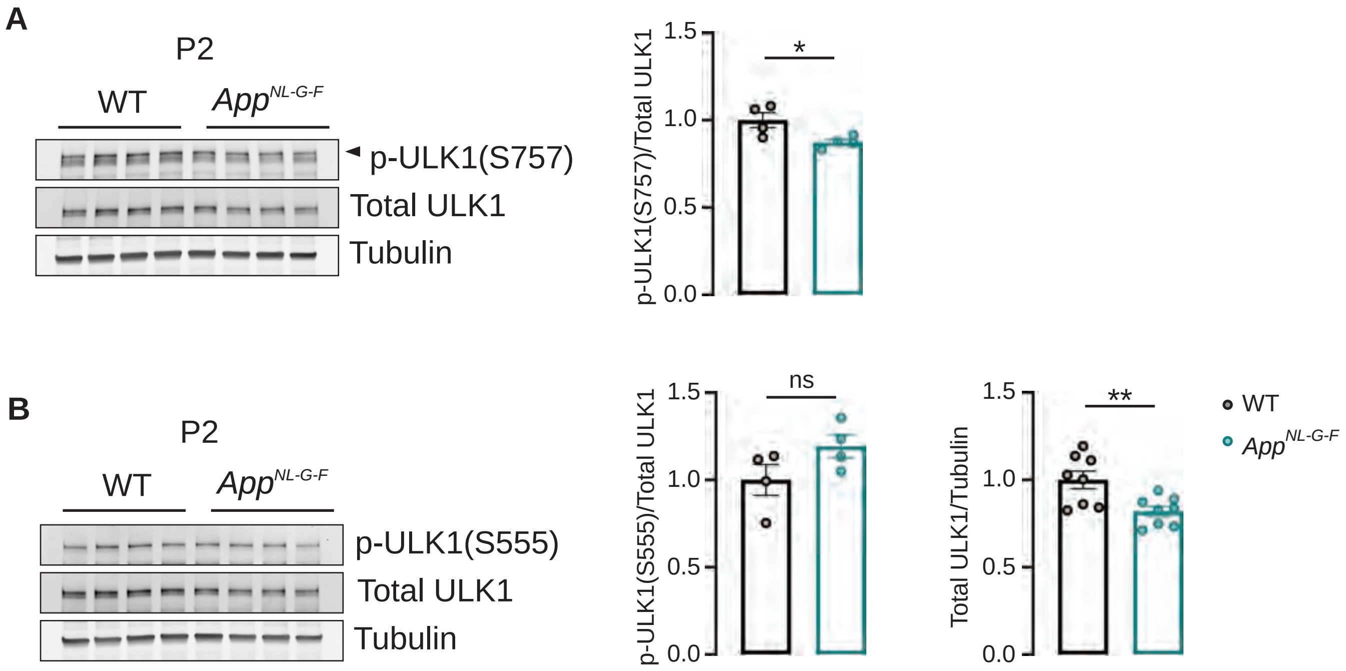
